# Unique brewing-relevant properties of a strain of *Saccharomyces jurei* isolated from ash (*Fraxinus excelsior*)

**DOI:** 10.1101/2021.01.11.426216

**Authors:** Mathias Hutzler, Maximilian Michel, Oliver Kunz, Tiina Kuusisto, Frederico Magalhães, Kristoffer Krogerus, Brian Gibson

## Abstract

The successful application of *Saccharomyces eubayanus* and *Saccharomyces paradoxus* in brewery fermentations has highlighted the potential of wild yeast for brewing, and prompted investigation into the application potential of other members of the genus. Here, we evaluate, for the first time, the brewing potential of *Saccharomyces jurei*. The newly isolated strain from an ash tree (*Fraxinus excelsior*) in Upper Bavaria, Germany, close to the river Isar, was used to ferment a 12°P wort at 15°C. Performance was compared directly with that of a reference lager strain (TUM 34/70) and the *S. eubayanus* type strain. Both wild yeast rapidly depleted simple sugars and thereafter exhibited a lag phase before maltose utilization. This phase lasted for 4 and 10 days for *S. eubayanus* and *S. jurei*, respectively. *S. eubayanus* utilized fully the available maltose but, consistent with previous reports, did not use maltotriose. *S. jurei*, in contrast, utilized approx. 50% of the maltotriose available, making this the first report of maltotriose utilization in a wild *Saccharomyces* species. Maltotriose use was directly related to alcohol yield with 5.5, 4.9, and 4.5 % ABV produced by *S. pastorianus, S. jurei* and *S. eubayanus*. Beers also differed with respect to aroma volatiles, with a high level (0.4 mg/L) of the apple/aniseed aroma ethyl hexanoate in *S. jurei* beers, while *S. eubayanus* beers had a high level of phenylethanol (100 mg/L). A trained panel rated all beers as being of high quality, but noted clear differences. A phenolic spice/clove note was prominent in *S. jurei* beer. This was less pronounced in the *S. eubayanus* beers, despite analytical levels of 4-vinylguaiacol being similar. Tropical fruit notes were pronounced in *S. jurei* beers, possibly resulting from the high level of ethyl hexanoate. Herein, we present the successful results of the first intentional application of S. *jurei* as a yeast for beer fermentation known to us and compare its fermentation performance to other species of the genus. Results indicate considerable potential for *S. jurei* application in brewing, with clear advantages compared to other wild *Saccharomyces* species.

## 1 Introduction

Yeasts of the genus *Saccharomyces* are by far the most prevalent fermentative microorganisms used in brewing. In addition to the commonly used yeasts *S. pastorianus* and *S. cerevisiae*, wild species of *Saccharomyces* such as *S. paradoxus* and *S. eubayanus* have recently been utilized intentionally for beer fermentations (Gibson, 2015; Osburn et al., 2016; Nikulin et al., 2020). The successful utilization of these non-domesticated species suggests that other species belonging to the genus could be profitably employed in beer production, and could facilitate differentiation of beers, or even creation of novel beer styles (Osburn et al., 2016). Alternative *Saccharomyces* species have the potential to introduce novel flavor profiles to beers and, in particular, to lager beers which are fermented at low temperatures (with the exception of *S. cerevisiae* the genus may be described as psychrophilic) (Magalhães et al., 2021). The potential benefits derived from wild yeasts are off-set by a number of characteristics that may be considered negative in the context of brewing. These include the production of phenolic flavor compounds (which typically lend a spice or smoke note to beer); poor flocculation, which hinders clarification; and inability to use all of the available sugars present in brewers wort, thereby limiting fermentation efficiency. Only *S. paradoxus* and *S. eubayanus* have been fully characterized with respect to brewing potential (Gibson et al. 2013; Nikulin et al. 2020, Mardones et al. 2020) and other species in the genus may prove to be more suitable for efficient production of flavorful beers.

Eight *Saccharomyces* species are currently recognized (Naseeb et al., 2017). These have been isolated from a range of habitats, but appear to be most prevalent in woodland environments. Oaks and other species within the *Fagaceae* family, for example, serve as a habitat for both *S. paradoxus* and *S. cerevisiae*. Sniegowski et al. have isolated several strains of *S. paradoxus* and *S. cerevisiae* from exudate, soil and bark associated with different trees belonging to the *Quercus* genus, while samples taken from poplar, maple and American beech trees did not yield *Saccharomyces* yeasts (Sniegowski et al., 2002). A clear preference of *Saccharomyces* for oak trees compared to trees outside the *Quercus* genus was also shown by Sampaio et al. (Sampaio and Gonçalves, 2008). Other *Saccharomyces* species, isolated from *Drosophila sp*. in Brazil, and from soil and decayed leaves in Japan, have been classified as *S. cariocanus, S. kudriavzevii* and *S. mikatae* respectively (Naumov et al., 2000) (though the former species is now often designated *S. paradoxus* (Liti et al., 2006)). *S. mikatae* has not been isolated from other locations to date suggesting a regional restriction of this species. *S. kudriavzevii* has also been isolated alongside other *Saccharomyces* species from oak bark and soil surrounding oak trees in Europe (Sampaio and Gonçalves, 2008). In 2008, a novel *Saccharomyces* species was isolated from trees in Western China. Three strains of the newly named *S. arboricola* were obtained from the bark of a *Quercus* sp. and a *Castanopsis* sp. tree, both belonging to the *Fagaceae* family (Wang and Bai, 2008). *S. eubayanus*, the cryotolerant co-parent of today’s widely used lager-brewing yeast *S. pastorianus*, was first isolated and identified on southern-beech trees (*Nothofagus spp*.) in Patagonia. Along with *S. eubayanus, S. uvarum* was observed in the same habitats indicating that these two cryotolerant species thrive in the cold climate of Patagonia (Libkind et al., 2011). These two species have later also been isolated in sympatry from trees in China, also including oaks (Bing et al., 2014) and Wisconsin, USA, on American beech and maple trees (Peris et al., 2014).

In 2017, Naseeb et al. first isolated and described *S. jurei* (NCYC 3947) from an oak tree (*Quercus robur*) in the French Pyrenees. They found a close relationship between *S. mikatae* and *S. jurei* through phylogenetic analysis and suggest shared evolutionary history of these two species (Naseeb et al., 2018). In a later study, (Alsammar et al., 2019) detected DNA homologous to that of *S. jurei* in natural habitats using high throughput sequencing of the ITS1 region specific to *Saccharomyces spp*. By sampling a variety of trees (oak, beech, spruce, larch and pine) at different altitudes in the Italian Alps, the soil surrounding the trees was scanned for evidence of the presence of members of the *Saccharomyces* genus. As expected, *S. cerevisiae* and *S. paradoxus* were abundant, but more interestingly, *S. mikatae* and *S. jurei* were identified in many samples as well as *S. eubayanus* and *S. kudriavzevii* (Alsammar et al., 2019). This did not yield any viable strains as only DNA was extracted from the soil. The apparent paucity of *S. jurei* in nature may be simply an artefact of the culturing methods used for isolation and enrichment, which may disadvantage some species relative to others. The use of wooden materials and tools in brewing has been widespread over centuries due to its relative ease of fabrication while metallic materials have only become the major material over the last century (Schnegg, 1921; 1922). The contact between wood harboring yeasts and wort containing fermentable sugars and nutrients may have be the source of yeast being used for brewing in the past or still today.

In this study, the potential of a *S. jurei* strain isolated from ash (*Fraxinus excelsior*) in Bavaria was investigated for its brewing potential. Traits studied included wort fermentation efficiency, sugar utilization, and beer flavor profile. Performance was compared to that of *S. eubayanus*, which has already been shown to be a capable brewing yeast, as well as the domesticated lager strain *Saccharomyces pastorianus* TUM 34/70.

## 2 Materials and Methods

### 2.1 Yeast strains

Strains used in this work are listed in Table 1. All strains with VTT codes were obtained from VTT culture collection (http://culturecollection.vtt.fi). The *S. pastorianus* strain TUM 34/70 and *S. jurei* strain TUM 629 were obtained from the TU Munich, Research Center Weihenstephan for Brewing and Food Quality (https://www.blq-weihenstephan.de/en/tum-yeast/yeast-and-bacteria/).

**Table 1.** Strains used in this study

| Species | Strain code and abbreviation | Other culture collection codes | Additional information |
| --- | --- | --- | --- |
| *S. cerevisiae* | VTT A-81062 (A62) |  | Ale yeast |
| *S. pastorianus* | TUM 34/70 |  | Lager yeast |
| *S. arboricola* | VTT C-15952<sup>T</sup> (C952) | CBS 10644<sup>T</sup> | Type strain |
| *S. eubayanus* | VTT C-12902<sup>T</sup> (C902) | CBS 12357<sup>T</sup> | Type strain |
| *S. kudriavzevii* | VTT C-15950<sup>T</sup> (C950) | CBS 8840<sup>T</sup>, ATCC MYA-4449<sup>T</sup>, NCYC 2889<sup>T</sup> | Type strain |
| *S. jurei* | VTT C-171003<sup>T</sup> (C1003) | CBS 14759<sup>T</sup>, NCYC 3947<sup>T</sup> | Type strain |
| *S. jurei* | TUM 629 |  | This study |
| *S. mikatae* | VTT C-15949<sup>T</sup> (C949) | CBS 8839<sup>T</sup>, NCYC 2888<sup>T</sup> | Type strain |
| *S. paradoxus* | VTT C-09850<sup>T</sup> (C850) | CBS 432<sup>T</sup>, NCYC 2600<sup>T</sup> | Type strain |

### 2.2 *Saccharomyces jurei* isolation, maintenance and microscopy

Samples of the bark of an ash tree (*Fraxinus excelsior*) located in the meadows of the Isar-river in Munich (Latitude 48.10931383333333 = 48 ° 6’ 33.53’’ N, Longitude 11.56397166666667 = 11° 33’ 50.298’’ E, 519.8 m above sea level) with attached moss were taken approximately 1 meter above ground with sterilized forceps and placed into sterile bags and stored for 2 days at 2 °C until processing. 1 g samples of the collected bark were put into a flask, sealed with a sterile plug, containing 50 mL of autoclaved wort at 12 °P (15 min at 121 °C), pH-value 5.3 (prepared from diluted wort concentrate (Döhler, Darmstadt Germany, original gravity approx. 60°) through addition of deionized water and tetracycline (tetracycline-hydrochloride, Carl Roth, Karlsruhe, Germany) at an application concentration of 50 mg/L. 0.25 ml of a saturated alcoholic biphenyl solution were pipetted onto the sterile plug to inhibit growth of mold. The flask containing wort and the sample was then incubated aerobically at 20 °C for 2 weeks.

An inoculation loop of the incubated medium was streaked on a Wallerstein Nutrient Agar plate (WLN-A, Oxoid GmbH, Wesel, Germany, pH 5,6, 50 mg/L tetracycline) and incubated anaerobically for one week at 20 °C. Single colonies were checked for cell morphology and purity of the colony with a microscope (Nikon, Düsseldorf, Germany) using dark field microscopy with a 1000-fold magnification. The examined colonies were then streaked onto a fresh WLN-A plate as described above. The colonies subsequently identified as *S. jurei* appeared small, round, and umbonate with a pale greenish-white coloration. A pure culture of *S. jurei* was streaked on a wort agar slant which was incubated for 3 days at 28 °C before storage at 2 °C. For strain maintenance recultivation was performed every month. For long term storage cryotubes in cryo-vials (Roti-Store yeast cryo vials, Carl Roth, Karlsruhe, Germany) at −80 °C were prepared according to the manufacturer’s directions.

### 2.3 Initial identification using species specific Saccharomyces Real-Time PCR assays and ITS, D1/D2 26S rDNA sequencing

Yeast DNA was isolated using a modified InstaGene Matrix (Biorad, Feldkirchen, Germany) protocol (Hutzler, 2009; Meier-Dörnberg et al., 2018). Yeast DNA of the single colony were analyzed using different species-specific *Saccharomyces* Real-Time PCR assays (Hutzler, 2009; Hutzler et al., 2015; J. P. Sampaio, 2017; Meier-Dörnberg et al., 2017b). Sequencing of ITS and D1/D2 26S rDNA loci was performed acccording to White et al., Kurtzman et al using modified protocols according to Hutzler (White, 1990; Kurtzman and Robnett, 1998; Kurtzman et al., 2003; Hutzler, 2009; 2010). Sequences were analyzed using NCBI Blast tool (NCBI) and DNAStar, MegAlign Software (DNASTAR, Inc., Madison, Wisconsin).

### 2.4 Whole-genome sequencing

Genomic DNA from strains *S. jurei* C1003 and TUM 629 was isolated using Qiagen 100/G Genomic-tips (Qiagen, The Netherlands). The DNA was sequenced at the Microbial Genome Sequencing Center (Pittsburgh, PA, USA). A 150-bp paired-end Illumina Nextera library was prepared, and sequencing was carried out with a NextSeq 550 instrument. The paired-end reads were trimmed and filtered with Trimmomatic (Bolger et al., 2014). Reads were aligned to the reference genome of *S. jurei* NCYC 3947^T^ (accession number GCA_900290405; (Naseeb et al., 2018)) using BWA-MEM (Li and Durbin, 2009). Variant analysis was performed on aligned reads using FreeBayes (Garrison and Marth, 2012). Prior to variant analysis, alignments were filtered to a minimum MAPQ of 50 with SAMtools (Li et al., 2009). Annotation and effect prediction of the variants were performed with SnpEff (Cingolani et al., 2012). The median coverage over 10,000-bp windows was calculated with mosdepth (Pedersen and Quinlan, 2017). Raw sequence reads have been deposited in the NCBI Short Read Archive under BioProject PRJNA681394.

### 2.5 Wort preparation

The wort for the fermentations was produced in the VTT pilot brewery. Milled Pilsner malt (Viking Malt, Lahti, Finland) was mashed in with local Espoo City water following an infusion mashing procedure (mashing-in at 48 °C; rests: 48 °C 30 min − 63 °C 30 min − 72 °C 30 min – 78 °C 10 min), mash was filtered with a Meura (Belgium) mash filter and boiled for 60 minutes with Magnum hop pellets (α-acid content 15%). The wort was hopped to achieve 40 IBU and the strength at knockout was 12 °Plato. The wort was collected hot (over 90 °C) in stainless steel kegs and stored at 0 °C before use. The concentrations of sugars in the wort were 52.5 g/L of maltose, 13.4 g/L of maltotriose, 11.6 g/L of glucose, and 2.7 g/L of fructose.

### 2.6 Fermentation trials and beer preparation

10 L-scale fermentations were carried out with *S. jurei* TUM 629, *S. eubayanus* C902, and *S. pastorianus* TUM 34/70. Yeasts were first propagated by transferring an inoculation loop of yeast from a YPD agar plate to 25 ml liquid YPD culture. The culture was incubated aerobically on a shaker for 24 h, before being transferred to 500 mL YPD. After aerobic incubation with agitation (120 rpm) on an orbital shaker for 3 days, the yeast suspension was centrifuged, a 20%-slurry (200g fresh yeast/L) was prepared in sterile Milli-Q-filtered water and yeast were inoculated into 1.5 L of 12 °P wort in a 2 L Schott-bottle capped with an airlock. After five days of static fermentation, the yeast was removed by centrifugation (4000 rpm; 5 min; 4 °C) and a 20% slurry was again prepared. Cell number was determined using the NucleoCounter YC-100 (ChemoMetec, Denmark) and cells were inoculated into 8 L of aerated (10 ppm dissolved oxygen) 12 °P wort in 10 L-volume, stainless-steel, cylindroconical vessels, to give a starting cell density of of 1 × 10^7^ cells/mL. Fermentations were conducted at 15 °C, and were monitored through regular sampling for assessment of wort pH, alcohol content and cell mass. After fermentation was complete, i.e. when minimal change in wort density was observed over consecutive days, the fresh beers were transferred from fermenters to kegs, matured for 7 days at 10 °C and stabilized seven days at 0 °C before depth filtration (Seitz EK, Pall Corporation, New York, NY, USA). Prior to bottling, the beers were carbonated to 5 g/L, and the bottled beers were stored at 0 °C.

### 2.7 Screening of *Saccharomyces* type strains for wort fermentation potential

Wort fermentation screening trials included six wild *Saccharomyces* species and two reference brewing strains: one ale (*S. cerevisiae*) and one lager (*S. pastorianus*) strain (Table 1). Prior to fermentation, an inoculation loop was used to transfer yeast from a stock YPD agar plate to the 50 ml liquid YPD medium in a 100 ml Erlenmeyer flask. The cultures were propagated at 20 °C on an orbital shaker (100 × g, Infors AG TR-125). After two days the yeast suspensions were centrifuged (10 min, 9000 × g and 4 °C) and a 20% (200 mg / ml) slurry was prepared for cell counting. The NucleoCounter YC-100 was used to calculate the cell count before the yeasts were transferred to the wort at a pitching rate of 1×10^7^ cells/mL. Cells were pitched according to cell number rather than mass due to the expected differences in cell size amongst the strains. Fermentations were carried out in 100 ml of the 12 °Plato all-malt wort and were conducted in 250 ml Erlenmeyer flasks, without agitation, at the typical lager brewing temperature of 15 °C for 40 days. Airlocks containing 2 mL of 85% glycerol were used to seal the flasks. Fermentation progress was monitored by measuring mass loss due to CO_2_ release. Fermentations were performed in duplicate. When fermentations were completed, samples were taken to assess alcohol content, yeast mass and viability.

### 2.8 Analytical methods

Alcohol content and pH-value of beer samples were determined using an Alcolyzer Plus with a DMA 5000 density meter and Xsample 122 sample changer (Anton-Paar GmbH, Ostfildern, Germany). Medium chain fatty acids and medium chain fatty acid esters were determined by gas chromatography with a flame ionization detector (GC-FID) with a 50 m 0.32 mm phenomenex-FFAP-0.25 µm column. The temperature protocol was 1 min 60 °C, 3 min 220 °C (5 °C/min), 8 min 240 °C (20 °C /min). Detector and injector temperatures were 250 °C and 200 °C, respectively. Fermentation by-products were determined using headspace GC-FID analysis according to Mitteleuropäische Brautechnische Analysenkommision method 2.21.1. Briefly, an INNOWAX cross-linked polyethylene-glycol 60 m × 0.32 mm 0.5 µm column was used. Temperatures of oven, detector and injector were 250 °C, 200 °C and 150 °C, respectively. Injection time was 4 s and analyzing time was 17 min. Turbo-Matrix 40 headspace parameters were: sample temperature, 60 °C; transfer temperature, 130 °C, needle temperature 120 °C. The time for GC-cycle was 22 min, thermosetting was 46 min, pressurization was 1 min and injection time was 0.03 min. Fermentable sugars and glycerol in beer were measured by HPLC. A 1.0 mL sample of wort or beer was filtered through Millipore membrane (pore size of 0.45 μm) filters and frozen (−20 °C). The samples were thawed and prepared for HPLC, which was used to determine concentrations of fermentable sugars (fructose, glucose, maltose and maltotriose) of wort and beers. A Waters 2695 Separation Module and Waters System Interphase Module liquid chromatograph coupled with a Waters 2414 differential refractometer (Waters Co., Milford, MA, USA) were used. An Aminex HPX-87H Organic Acid Analysis Column (300 × 7.8 mm, Bio-Rad, USA) was equilibrated with 5 mM sulphuric acid (H_2_SO_4_) (Titrisol, Merck, Germany) in water at 55 °C. The samples were eluted with 5 mM H_2_SO_4_ in water at a 0.3 mL/min flow rate.

### 2.9 Maltotriose transport assays

For maltotriose uptake measurement, the yeast strains were grown at 20 °C in liquid YP medium containing maltose (4% w/v) or maltotriose (4% w/v) to an OD600 nm between 4 and 8. The cells were harvested by centrifugation (10 min, 5000 rpm, 0°C), washed with ice-cold water and 0.1 M tartrate-Tris (pH 4.2) and re-suspended in the same buffer to a concentration of 200 mg/mL fresh yeast. Zero-trans rates of [U-^14^C]-maltotriose uptake were measured at 20 °C essentially as described by (Lucero et al., 1997). Briefly, aliquots of 40 µl of yeast suspension were added to 20 µl of 15 mM labeled maltotriose (for a final concentration of 5 mM [U-^14^C]-maltotriose) and incubated for 60 s at 20 °C. The reaction was stopped with the addition of 5 ml ice-cold water. The suspension was quickly filtered and washed with an additional 5 ml of ice-cold water. The filter was submerged in 3.5 ml of Optiphase HiSafe 3 scintillation cocktail (Perkin Elmer, MA, USA) and the radioactivity measured in a Perkin Elmer Tri-carb 2810 TR scintillation counter. [U-^14^C]-maltotriose (ARC 627) was obtained from American Radiolabeled Chemicals (St. Louis, MO, USA) and re-purified before use as described by (Dietvorst et al., 2005). Maltose (minimum purity, 99%) and maltotriose (minimum purity, 95%) were from Sigma-Aldrich (St. Louis, MO).

### 2.10 Sensory analysis

All beer samples were tasted and judged by a trained sensory panel of 10 panelists certified by the Deutsche Landwirtschafts-Gesellschaft (DLG). Single tasting was performed in a dedicated tasting room (single tasting chambers, white-colored room, no distracting influences, and brown glasses with three-digit number labels) to exclude all external misleading factors. The main flavor impressions were determined at a range from 1 (almost no perception) to 10 (very intense perception). Flavor impressions were chosen according to Meier-Dörnberg et al. (Meier-Dörnberg et al., 2017a). In addition, a tasting was performed under the same circumstances with the DLG scheme, in which the beer is judged by its aroma, taste, carbonation, body and bitterness in a range of 1 to 5, 1 being the lowest value (negative) and 5 being the highest value (positive).

## 3 Results

*S. jurei* TUM 629 was isolated from a piece of bark of an ash tree (*Fraxinus excelsior*) close to the river Isar in Munich, Bavaria, Germany. Single colonies of *S. jurei* TUM 629 exhibited a homogenous morphology (Figure 1). On WLN-Agar, colonies appear white with a pale green center, and exhibit a well-defined edge and a distinct umbonate morphology. On wort-agar, the colonies are round and white becoming more transparent towards the edge. The colonies of *S. jurei* TUM 629 differed in colony morphology from the colonies of *S. eubayanus* C902 and *S. pastorianus* TUM 34/70 (both of which were later used for the brewing trials as reference strains). As both agars are standard tools within brewing microbiology they may serve as useful preliminary differentiation tools for *S. jurei* monitoring.

**Figure 1.**
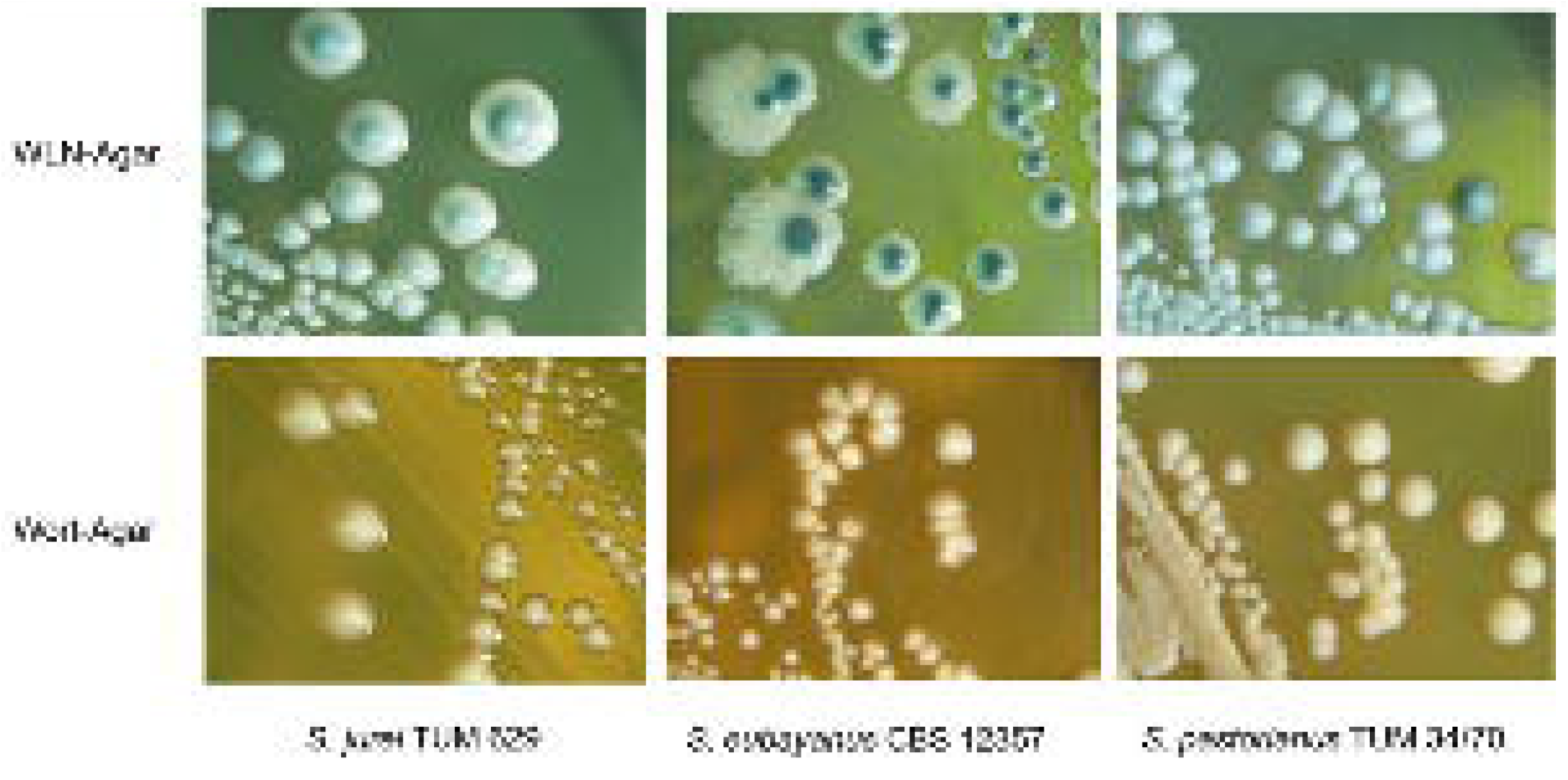
Colony morphologies of *S. jurei* TUM 629, *S. eubayanus* C902, *S. pastorianus* TUM 34/70 on WLN-Agar (Wallerstein-Nutrient-Agar) and Wort-Agar.

The cell morphology of *S. jurei* in brewer’s wort (12 °P pale barley malt wort) is round with single cell-budding of round daughter-cells (dark-field microscopy with scale in Figure S1). Cell diameter is between 4 and 8 µm. Vacuoles and cell organelle structures could be observed in the dark field microscopic picture. Cell morphology of *S. jurei* TUM 629 in wort is different from the cell morphologies of *S. pastorianus* lager strains and from some *S. cerevisiae* brewing strains (e.g. wheat beer strains with larger cell diameter and multilateral budding and star cluster formation).

The ITS1-5.8-ITS2 rDNA and D1/D2 units of the 26S rDNA of *S. jurei* TUM 629 were sequenced and NCBI Blast comparison was carried out (sequences in supplementary data). D1/D2 26S rDNA showed 100 % sequence identity and ITS1-5.8-ITS2 rDNA 99.38 % sequence identity to *S. jurei* type strain NCYC 3947^T^ (data not shown). Figure 2 gives a clear indication of species identity of TUM 629 by showing the median coverage of 10 kbp windows of seqeuencing reads to a reference genome of 8 *Saccharomyces* species. A phylogenetic tree based on the Clustal W Alignment of the two *S. jurei* sequences (TUM 629 and NCYC 3947^T^) along *S. pastorianus* and *S. eubayanus* further supporting the species identity can be found in supplementary material (Figures S2 and S3).

**Figure 2.**
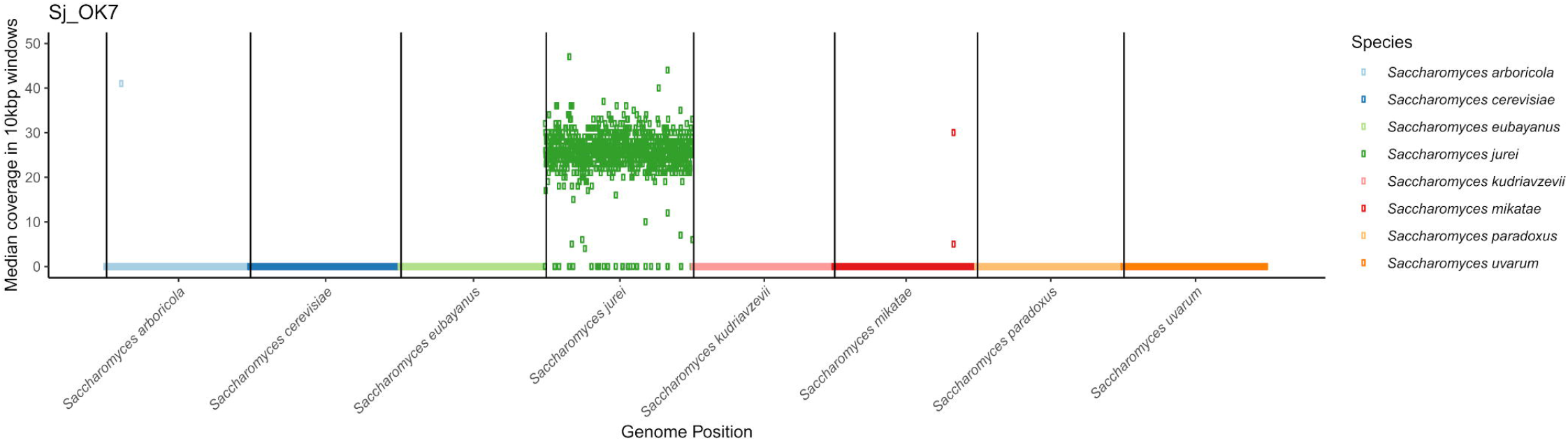
The median coverage in 10 kbp windows of sequencing reads from *S. jurei* TUM 629 aligned to a concatenated reference genome consisting of 8 species in the *Saccharomyces* genus. Reads align exclusively to *S. jurei*, except for two small regions in the sub-telomeric regions of *S. arboricola* chromosome 4 and *S. mikatae* chromosome 14. Results were visualized in R using modified scripts from sppIDer (Langdon et al., 2018).

### 3.1 Genome sequencing and analysis

To confirm the species-level identification of *S. jurei* TUM 629, 150 bp paired-end Illumina whole genome sequencing was carried out. *S. jurei* TUM 629 and VTT C-171003 (= C1003 = NCYC 3947^T^) were sequenced to an average coverage of 29× and 27×, respectively. A total of 9690 single nucleotide polymorphisms (SNPs) were detected in TUM 629 compared to the reference genome of *S. jurei* C1003 (accession number GCA_900290405; (Naseeb et al., 2018)). Of these SNPs, only 303 were heterozygous, suggesting that TUM 629 is a homozygous diploid. The SNPs in TUM 629 indicate a sequence divergence of about 0.1% relative to the type strain, which was originally isolated in south-eastern France (Naseeb et al., 2017). Sequencing coverage was even across the whole reference genome, suggesting that TUM 629 is euploid (Figure S4, Figure S5 showing the coverage for *S. jurei* C1003). When sequencing reads were aligned to a concatenated reference genome of all *Saccharomyces* species, reads mapped exclusively to *S. jurei*, indicating that *S. jurei* TUM 629 is not an interspecies hybrid (Figure 2). As expected, only 222 SNPs (218 of which were heterozygous) were detected in the re-sequenced *S. jurei* type strain compared to the same reference genome. Genetic analysis allowed identification of genes potentially encoding maltose and maltotriose transporter proteins (Table S1). Three genes ranging from 82.2% to 84.4% identity with *S. cerevisiae MAL31* and one with 82.6% identity with *S. cerevisiae MAL11* were identified. Additionally, 3 genes were found with 78.9% to 84% identity to *S. cerevisiae IMA5*, encoding an extracellular α-glucosidase.

### 3.2 Wort fermentation

Wort fermentation was carried out to assess the brewing-relevant properties of *S. jurei* TUM 629. Results were compared with those of the type strain of *S. eubayanus* (C902), and the *S. pastorianus* lager strain TUM 34/70 (Figure 3). An initially rapid fermentation rate in the first two days was followed by a period of relative inactivity, which lasted for approximately 14 days. After this time alcohol level increased steadily for approx. 30 days, reaching a value of 4.9 % (v/v) at 39 days. *S. eubayanus* likewise exhibited rapid fermentation in the first two days, and this was also followed by a lag phase. In contrast to the *S. jurei* fermentation, this period of inactivity only lasted until 7 days after inoculation before the fermentation rate increased. Fermentation was mostly complete after 18 days, reaching a maximum of 4.4 % alcohol. As expected, fermentation with the lager strain was rapid, with no evidence of a lag phase at any stage. 5.5 % (v/v) alcohol was achieved after 9 days with this strain. Analysis of residual sugars indicated that maltotriose consumption was a determining factor for the extent of fermentation (Figure 4). Approximately 50 % of the available maltotriose was consumed by *S. jurei*. The corresponding depletion in the beers fermented with the lager strain was 85 %. *S. eubayanus*, as expected, had no effect on maltotriose concentration. Maltose was completely consumed by the lager strain and by *S. eubayanus*, though interestingly a small portion (3.6 g/L of initially 52.5 g/L) remained in the *S. jurei beer*. An increase in glycerol after fermentation was observed for all strains. This was highest in the wild yeast strains both of which produced 2.9 g/L of glycerol while the corresponding concentration in the lager strain beer was 1.6 g/L.

**Figure 3.**
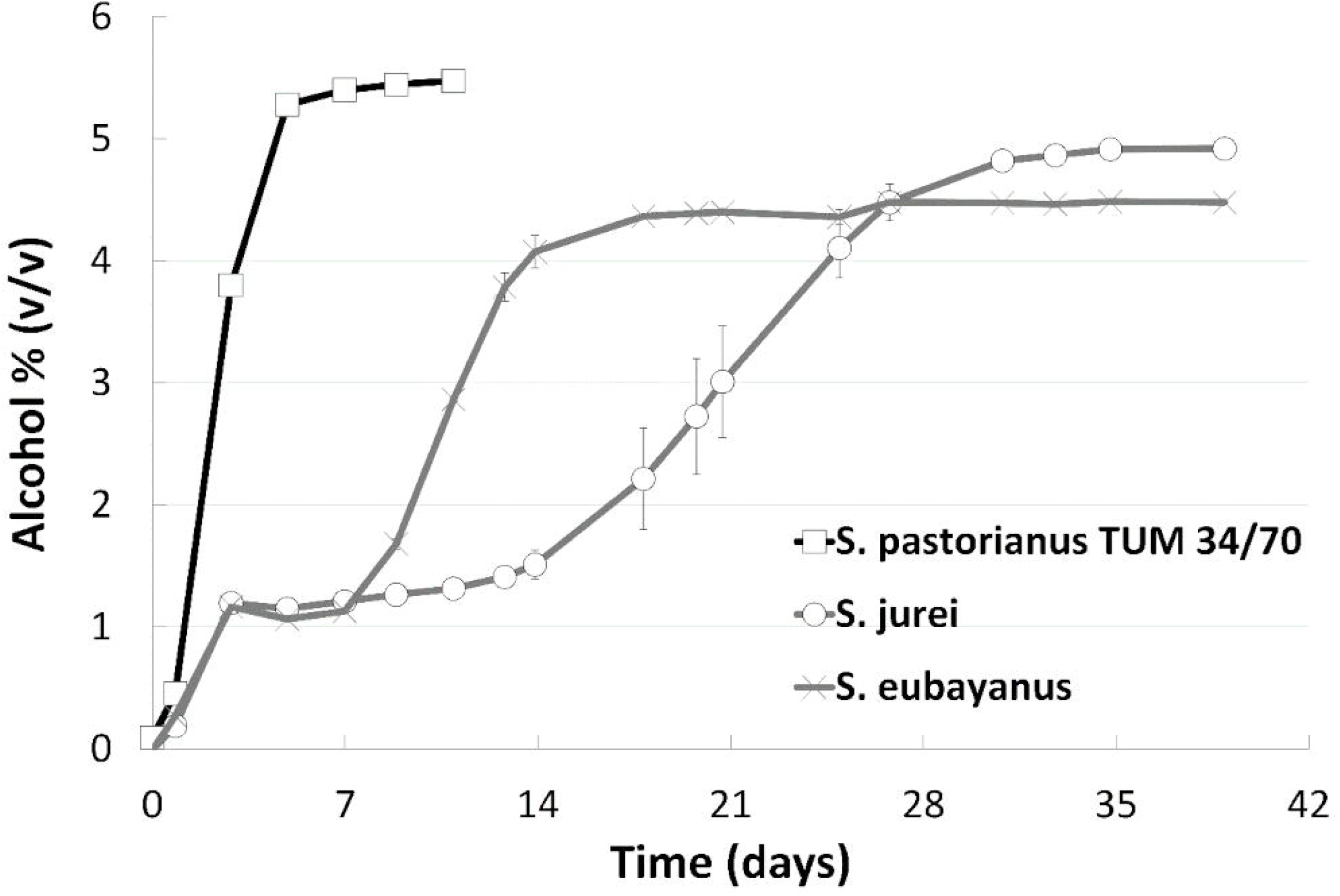
Fermentation progress as monitored by alcohol level (v/v) at 15 °C during 39 days of fermentation of an all-malt 12 °P wort. Strains include one lager reference strain (TUM 34/70), as well as the German strain of *S. jurei* (TUM 629), and the type strain of *S. eubayanus* (C902) Values are means from duplicate fermentations and error bars indicate range.

**Figure 4.**
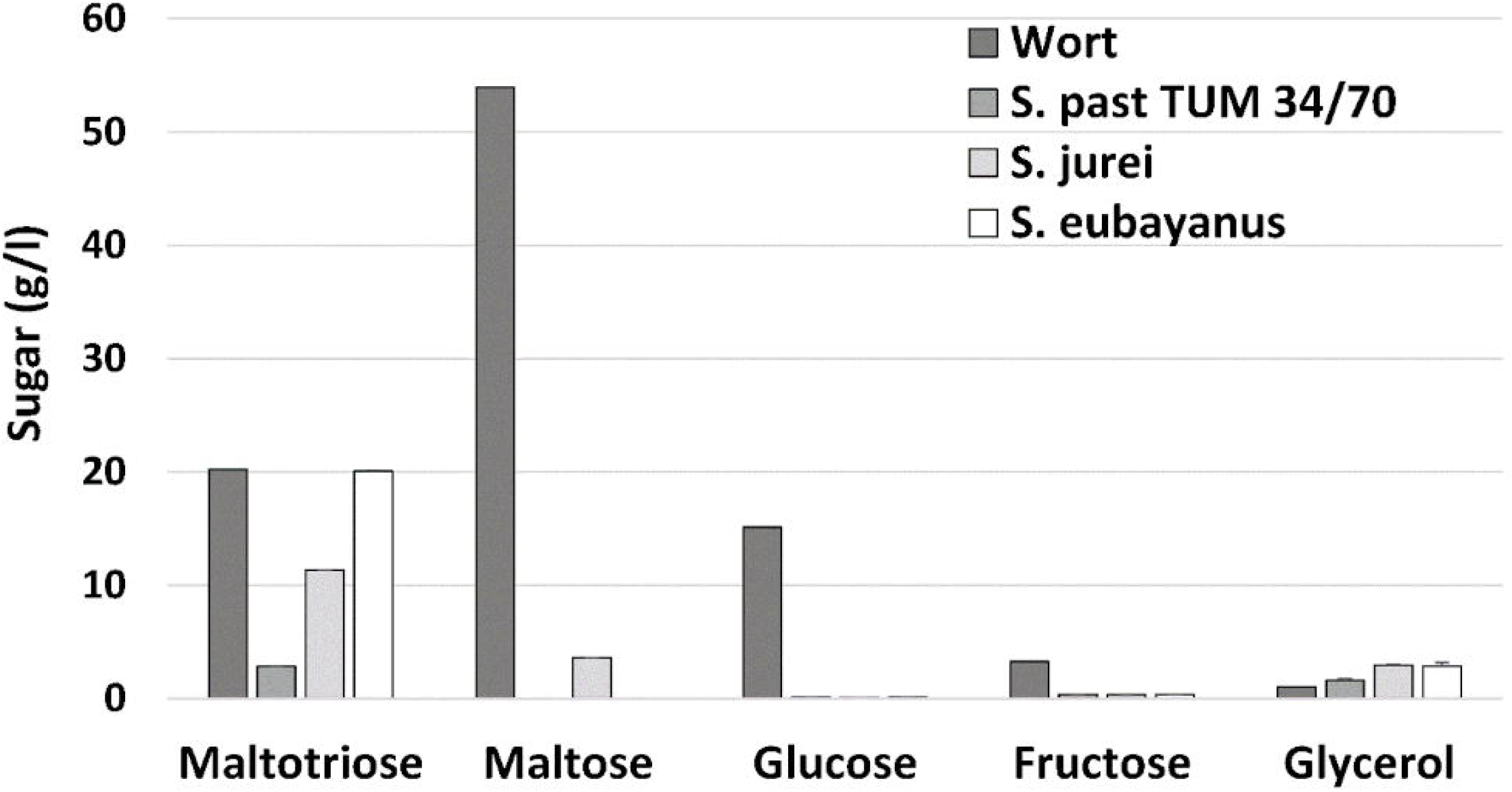
Residual sugars and glycerol in all-malt 12 °P wort, and beer after fermentation at 15 °C for up to 39 days. Strains include one lager reference strain (TUM 34/70), as well as the German strain of *S. jurei* (TUM 629), and the type strain of *S. eubayanus* (C902). Values are means from duplicate fermentations and error bars where visible indicate range.

### 3.3 Maltotriose uptake

To assess the yeast’s ability to take up maltotriose directly from the medium, uptake activities were quantified using radiolabeled maltotriose. When using maltose as propagation substrate, both *S. jurei* strains and *S. mikatae* showed maltotriose uptake activities ≤0.5 µmol min^-1^ g^-1^ DY^- 1^ indicating the absence of active maltotriose transporters in the plasma membrane (Table 2). An uptake activity ≤0.5 μmol min^−1^ g^−1^ DY is considered negligible. *S. eubayanus* C902 showed an activity level slightly above 0.5 μmol min^−1^ g^−1^ DY, although it is known that this strain lacks any capacity to transport maltotriose (Gibson et al., 2013; Magalhães et al., 2016). *S. pastorianus* was the only species showing enough maltotriose uptake activity to ensure its consumption from the wort. The fermentation data however showed that *S. jurei* can consume maltotriose (Figure 4). If the maltotriose is taken up by the yeast cells then the transporters are either susceptible to repression by maltose or its expression requires induction by maltotriose. To validate this hypothesis the maltotriose uptake activity was also measured in cells propagated in YP medium with maltotriose as sole carbon source. Growth on maltotriose was slow, and it took 6 days to reach an OD600 value greater than 4, however, after this maltotriose exposure, the cells could grow much faster when transferred to fresh medium containing maltotriose as sole carbon source (data not shown). Uptake activity from cells grown on maltotriose confirmed that there is an active transmembrane transport occurring in *S. jurei*. This mechanism appears to require prolonged exposure to maltotriose, or absence of other sugars.

**Table 2.** Zero-trans rates of maltotriose uptake activity (μmol min^−1^ g^−1^ DY) of the strains in study, measured at 20°C. Prior to assessment of activity, strains were propagated in YP medium supplemented with either maltose or maltotriose. Values are means of two independent assays (± standard deviation).

|  |  |
| --- | --- |
|  | Propagation media |

| Strains | YP + Maltose | YP + Maltotriose |
| --- | --- | --- |
| *S. jurei* C1003 | 0.5 ± 0.11 | 6.2 ± 0.93 |
| *S. jurei* TUM 629 | 0.5 ± 0.10 | 8.0 ± 0.07 |
| *S. eubayanus* C902 | 0.8 ± 0.03 | n.q. |
| *S. mikatae* C949 | 0.1 ± 0.05 | n.q. |
| *S. pastorianus* TUM 34/70 | 7.4 ± 0.81 | n.q. |

n.q. - not quantified

### 3.4 Analytical and sensory aroma profile

The beers produced by the three different yeast strains *S. pastorianus* TUM 34/70, *S. eubayanus* C902 and the newly isolated *S. jurei* TUM 629 showed significantly different sensory and analytical aroma profiles (Tables 3 and 4). The sample fermented by *S. pastorianus* TUM 34/70 had the overall highest concentration of esters (Table 3). Of all esters detected in this sample, ethyl acetate was found at the highest concentration at 42.1 mg/L. The overall highest concentration of 3-methylbutylacetate (iso-amyl acetate) was 3.6 mg/L, and was also found in the *S. pastorianus* beer. Ethyl hexanoate concentration was four times higher in the sample fermented with *S. jurei* TUM 629 (0.40 mg/L) compared to the samples fermented with *S. pastorianus* (0.13 mg/L) and *S. eubayanus* CBS 12537 (0.11 mg/L). Ethyl hexanoate (apple, fruity flavor) has a flavor threshold in beer of 0.23 mg/L according to Meilgaard (Meilgaard, 1975). The sample fermented by *S. jurei* showed a noticeable apple flavor as shown by the tasting results in Figure 5, which correlates well with the aforementioned analytical findings. Overall, *S. jurei* produced higher amounts of medium chain fatty esters and relatively high amounts of higher alcohols in comparison to the other two yeast strains but was outperformed in the production of acetate esters by the *S. pastorianus* strain, and in higher alcohol production by the *S. eubayanus* strain (Tables 3 and 4). *S. eubayanus* produced a typically high amount of phenyl ethanol (121.4 mg/L) in comparison to the other two yeast strains (Gibson et al., 2013). All samples showed diacetyl levels below 0.1 mg/L and no 2,3-pentanedione (data not shown).

**Table 3.**
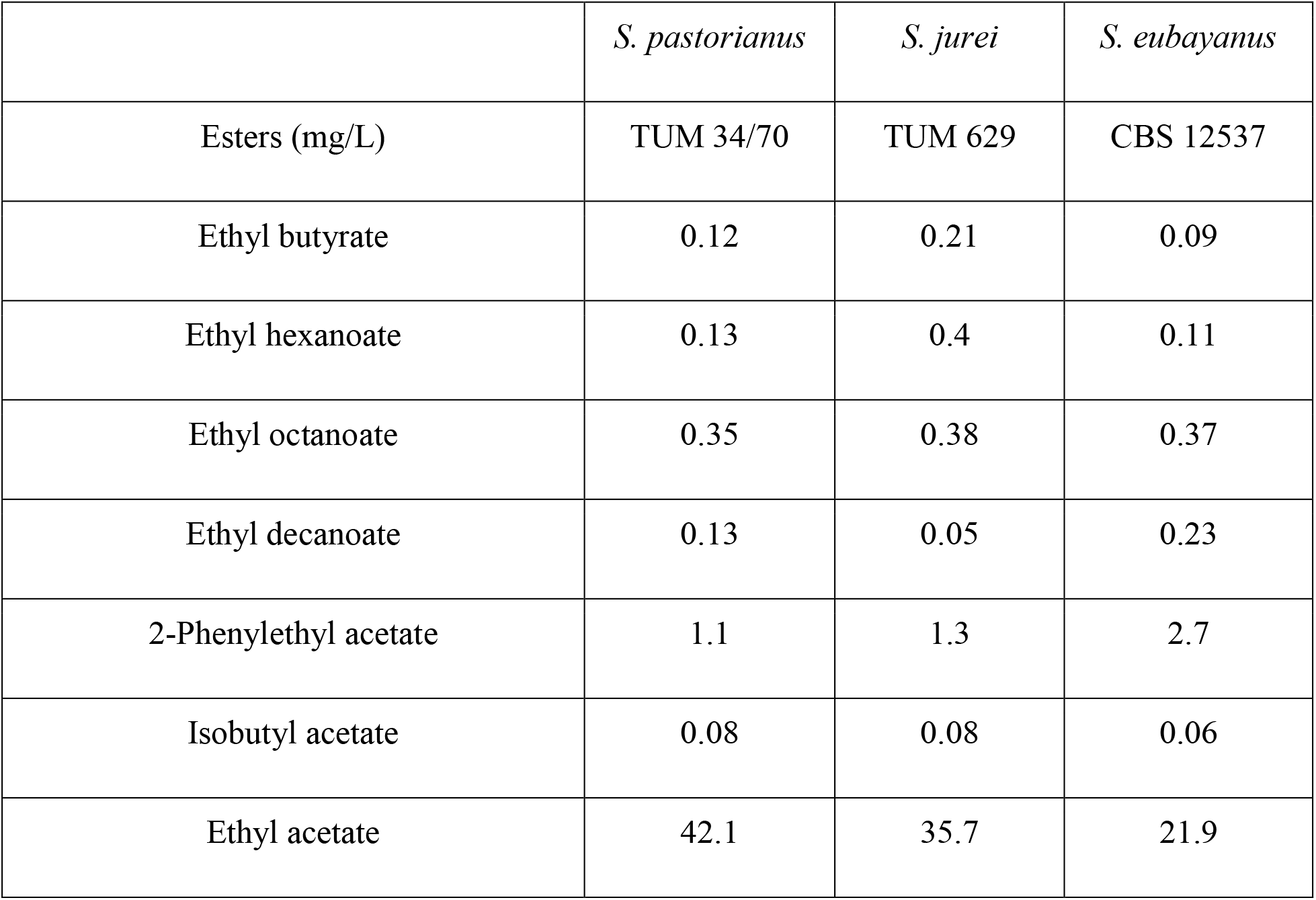
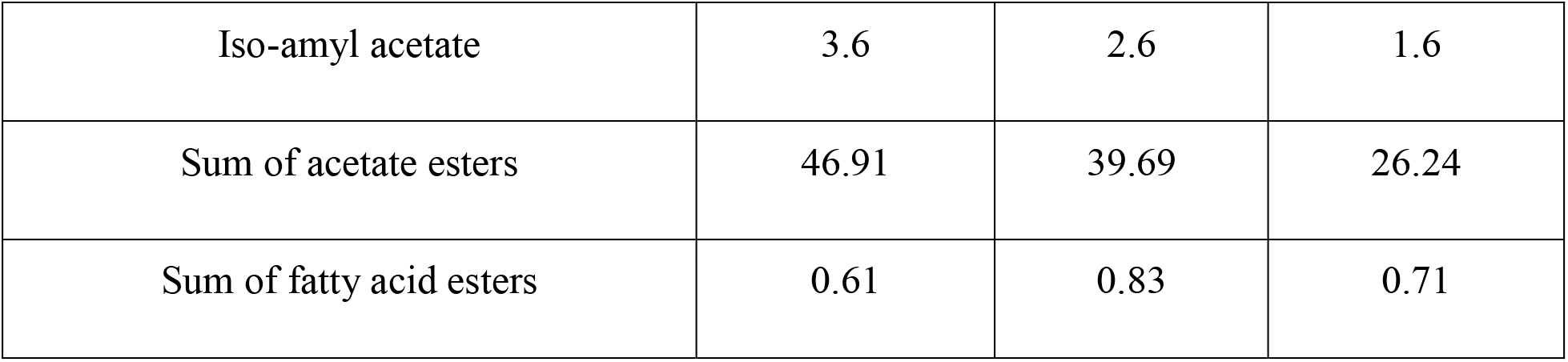
Esters detected in the three different beer samples fermented by *S. pastorianus* (TUM 34/70), *S. jurei* (TUM 629) and *S. eubayanus* (CBS 12537) at a fermentation temperature of 15 °C and an original gravity °12 P.

|  | *S. pastorianus* | *S. jurei* | *S. eubayanus* |
| --- | --- | --- | --- |
| Esters (mg/L) | TUM 34/70 | TUM 629 | CBS 12537 |
| Ethyl butyrate | 0.12 | 0.21 | 0.09 |
| Ethyl hexanoate | 0.13 | 0.4 | 0.11 |
| Ethyl octanoate | 0.35 | 0.38 | 0.37 |
| Ethyl decanoate | 0.13 | 0.05 | 0.23 |
| 2-Phenylethyl acetate | 1.1 | 1.3 | 2.7 |
| Isobutyl acetate | 0.08 | 0.08 | 0.06 |
| Ethyl acetate | 42.1 | 35.7 | 21.9 |
| Iso-amyl acetate | 3.6 | 2.6 | 1.6 |
| Sum of acetate esters | 46.91 | 39.69 | 26.24 |
| Sum of fatty acid esters | 0.61 | 0.83 | 0.71 |

**Table 4.**
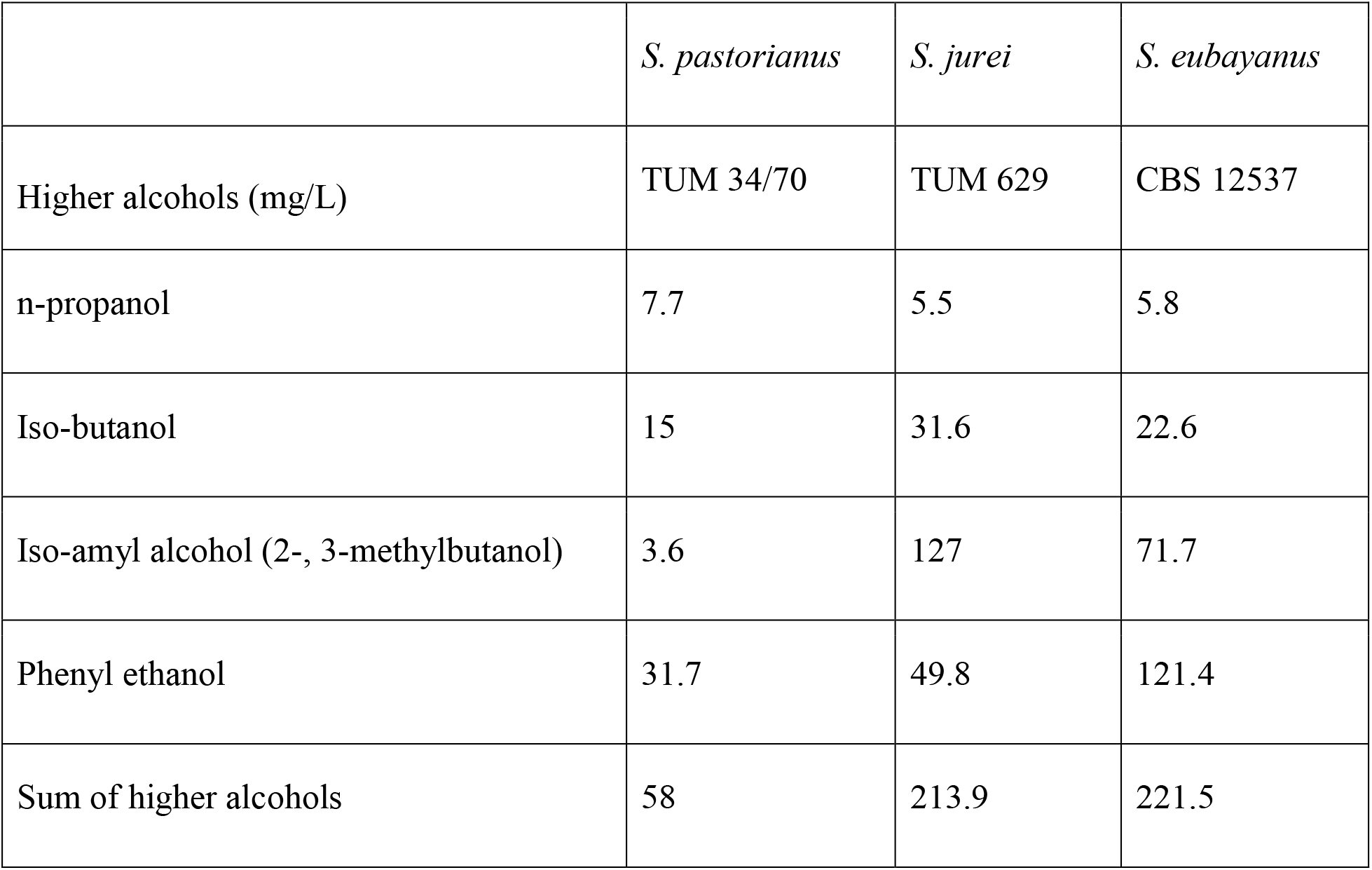
Higher alcohols detected in the three different beer samples fermented by *S. pastorianus* (TUM 34/70), *S. jurei* (TUM 629) and *S. eubayanus* (CBS 12537) at a fermentation temperature of 15 °C and an original gravity °12 P.

|  | *S. pastorianus* | *S. jurei* | *S. eubayanus* |
| --- | --- | --- | --- |
| Higher alcohols (mg/L) | TUM 34/70 | TUM 629 | CBS 12537 |
| n-propanol | 7.7 | 5.5 | 5.8 |
| Iso-butanol | 15 | 31.6 | 22.6 |
| Iso-amyl alcohol (2-, 3-methylbutanol) | 3.6 | 127 | 71.7 |
| Phenyl ethanol | 31.7 | 49.8 | 121.4 |
| Sum of higher alcohols | 58 | 213.9 | 221.5 |

**Figure 5.**
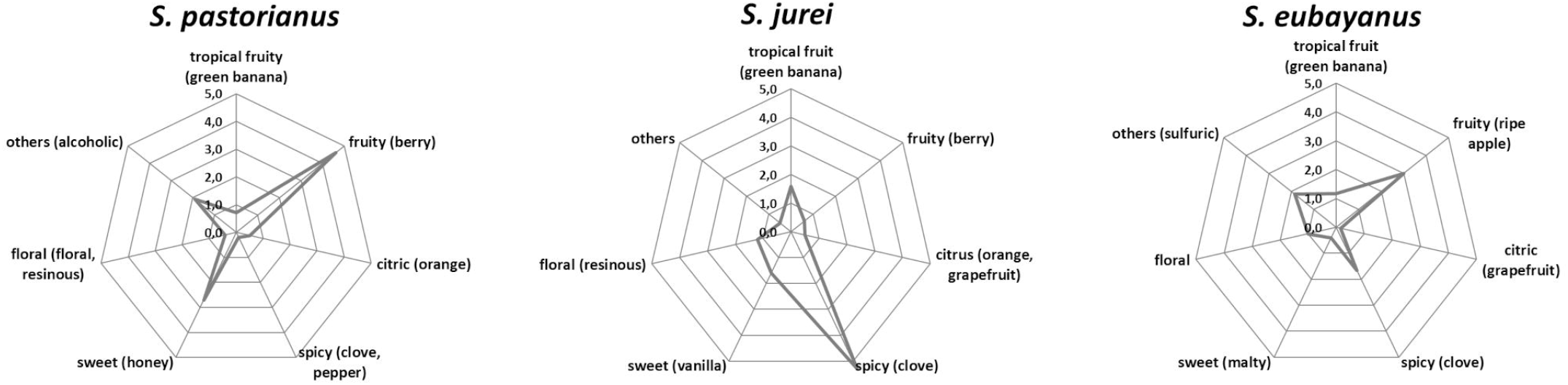
Tasting results of the beer samples fermented by *S. pastorianus* TUM 34/70, *S. jurei* TUM 629, and *S. eubayanus* at 15 °C fermentation temperature and an original gravity of 12 °P.

The trained panel rated all beers as high quality in the DLG scheme and found no significant difference of the purity of aroma and taste, and the quality of carbonation, body and bitterness among the three samples (data not shown). However, they noted clear differences in the descriptive analysis (Figure 5). A phenolic spice/clove note was prominent in the sample fermented by the *S. jurei* strain TUM 629. Further as mentioned above, a significant apple flavor was detected, which can be related to the considerably higher amount of ethyl hexanoate (Table 3). The clove-like flavor was less pronounced in the sample fermented by S. *eubayanus*, despite the analytical levels of 4-VG being similar (*S. eubayanus*: 0.16 mg/L, *S. jurei*: 0.15 mg/L). The sample fermented by *S. pastorianus* TUM 34/70 was described as very fruity and berry like, which can partly be explained by generally higher acetate ester concentration (46.91 mg/L) in the sample in comparison to the other two samples (*S. eubayanus* 26.24 mg/L and *S. jurei* 39.69 mg/L) (Table 3). Results indicated considerable potential for *S. jurei* application in brewing, and prompted a comparative study including the type strain of *S. jurei* and a number of other *Saccharomyces* type strains

### 3.5 Screening of *Saccharomyces* type strains for wort fermentation potential

Trial fermentations conducted with 12 °P wort at 15 °C revealed clear differences between the strains in terms of wort fermentation potential (Figure 6). As expected, the two reference strains fermented rapidly, with no evidence of an extended lag phase. Alcohol yield was good for both strains, 5.8 % ABV for *S. pastorianus* and 6.1 % ABV for *S. cerevisiae* (Figure 6). The greater efficiency of the A62 ale strain is due to a strong maltotriose fermentation capacity which has also been observed in previous studies (Krogerus et al., 2018).

**Figure 6.**
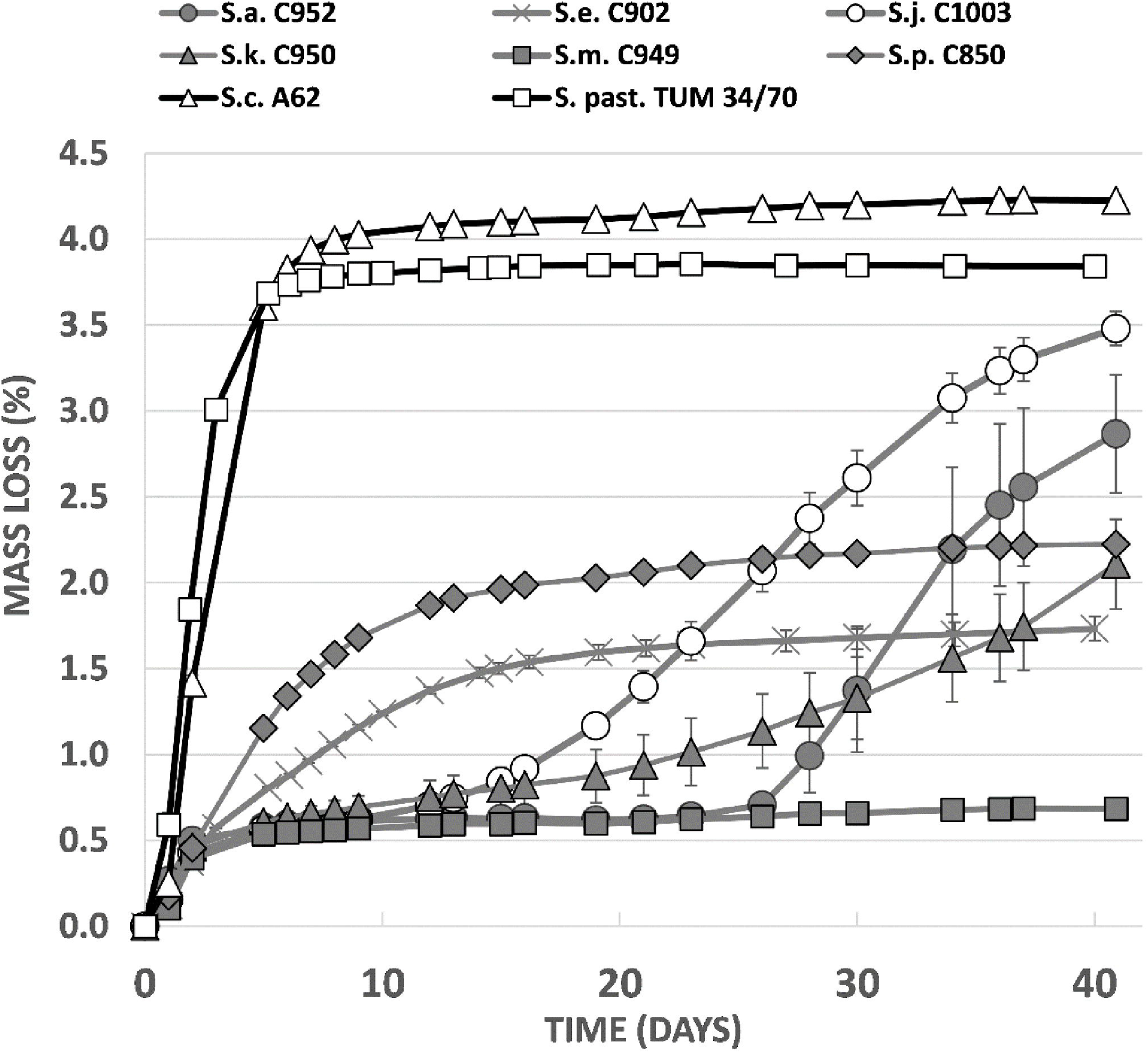
Fermentation performance (measured as loss of mass of the fermentor) in comparison between *Saccharomyces* type strains in 12 °P all-malt wort at 15 °C.

Fermentation characteristics of the *Saccharomyces* type strains were highly variable (Figure 6). All strains exhibited a relatively rapid fermentation in the first two days after inoculation, presumably due to utilization of the simple sugars present in the wort. *S. mikatae* fermentation appeared to cease after this initial period, while *S. arboricola*, which had an identical fermentation profile until 26 days after inoculation, appeared to start fermenting and was still actively fermenting after 41 days when it had produced an ABV level of 4.5 %. *S. kudriavzevii* fermentations were likewise characterized by an initially slow fermentation rate, which increased over time, giving a final value of 3.6 % ABV. *S. paradoxus* and *S. eubayanus*, did not exhibit any lag phase in fermentation, but overall fermentation efficiency was limited with the strains achieving 3.6 and 3.3 % ABV after 41 days (Figure 7). *S. jurei*, despite an initially low fermentation rate in the first two weeks of fermentation was able to achieve an ABV of 5.3%, a value considerably higher than those found in the other wild yeast beers.

**Figure 7.**
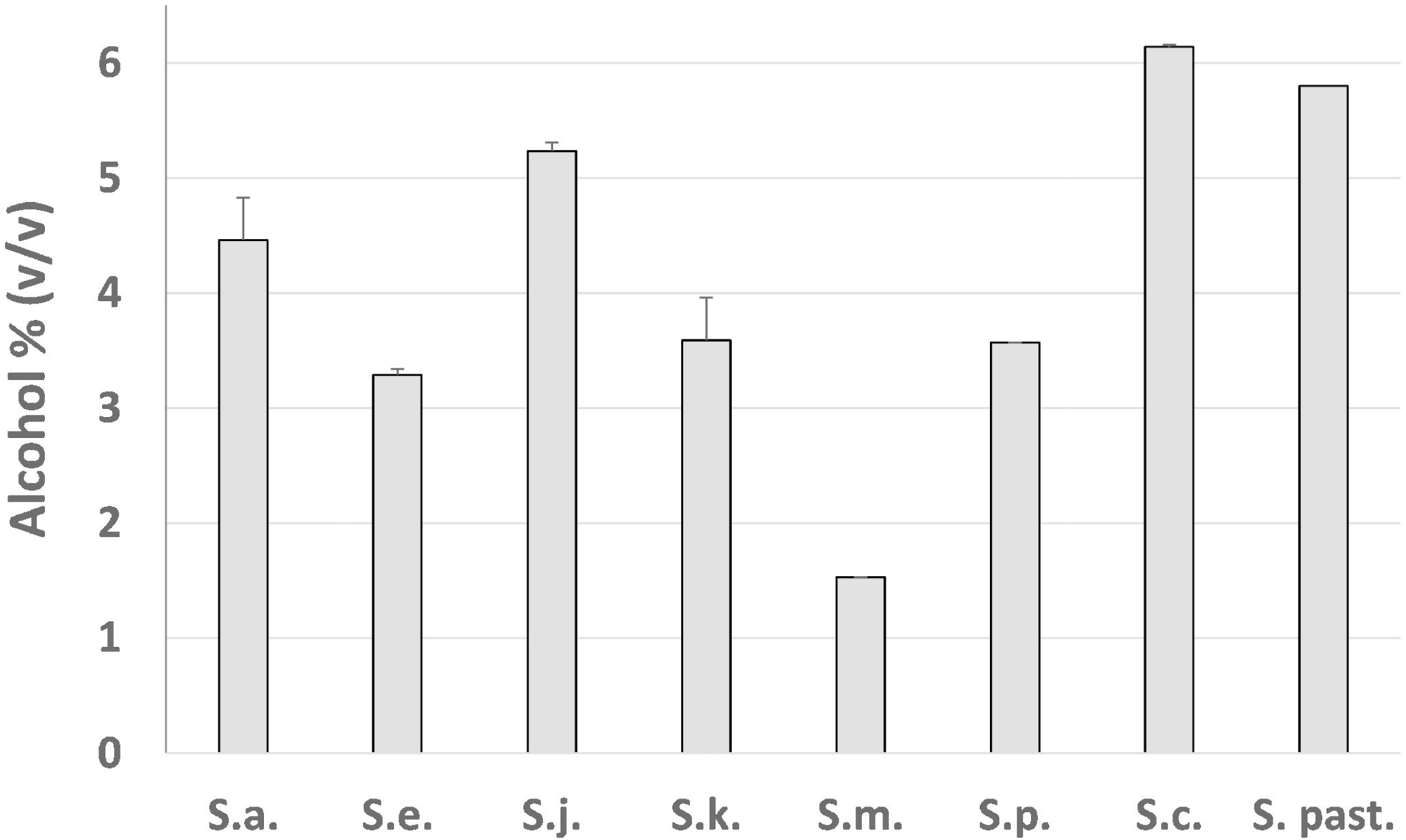
Final alcohol content of fermentations conducted in 12 °P brewer’s wort at 15 °C with *Saccharomyces* type strains.

## 4 Discussion

With the exception of recent reports on the brewing potential of two wild *Saccharomyces* species, *S. eubayanus* and *S. paradoxus* (Gibson et al., 2013; Cubillos et al., 2019; Nikulin et al., 2020), little is known about the brewing potential of the wild species within the *Saccharomyces* genus. Wild *Saccharomyces* species are not typically isolated from brewery fermentation environments, and when encountered in brewing, or other fermentation environments, typically occur in the form of interspecies hybrids. This has been seen for example with *S. kudriavzevii*, which occurs in a *S. cerevisiae* × *S. kudriavzevii* hybrid form in Belgian ales (Gallone et al., 2019), but is not otherwise isolated from brewing systems. Strains of *S. cerevisiae var. diastaticus* are common brewery contaminants (Meier-Dörnberg et al., 2018), but their designation as ‘wild’ species is questionable considering they group phylogenetically with one of the major ale yeast clades (Krogerus and Gibson, 2020; Pontes et al., 2020). The absence of wild yeast species in fermentation environments is indicative of deficiencies in performance. However, the successful application of *S. eubayanus* in brewing (Gibson et al., 2017) has demonstrated how such strains, when handled appropriately, may act as efficient fermenters, and can support product differentiation. Creative utilization of wild species may help brewers meet the consumer demand for beers with novel flavor profiles and interest in engaging background narratives.

The ability of *S. jurei* to utilize wort maltotriose was surprising given that this trait has not been observed previously in wild *Saccharomyces* yeasts and is generally considered to be a trait associated only with domesticated yeasts (Gallone et al., 2018). There are two mechanisms by which *Saccharomyces* yeast may utilize maltotriose (Krogerus et al., 2019). In the first, maltotriose is taken up directly across the cell membrane and hydrolyzed to glucose monomers through the action of an intracellular maltase. Active uptake is mediated by transporters such as AGT1 and MTT1, which are also responsible for the uptake of maltose (Dietvorst et al., 2005; Vidgren and Londesborough, 2012). Despite the absence of the property in wild species, maltotriose utilization appears to be evolvable, with two separate studies demonstrating the creation of maltotriose transporters from existing maltose transporters via a series of recombination events (Baker and Hittinger, 2019; Brouwers et al., 2019). A second mechanism for maltotriose utilization involves extracellular degradation of maltotriose by ‘diastatic’ strains of yeast. This is seen in some *S. cerevisiae* strains belonging, in particular, to the Beer 2 group (Gallone et al., 2016). In many of these strains, glucoamylase activity is responsible for the liberation of glucose and maltose from maltotriose, as well as dextrins and soluble starch, and obviates the requirement for transmembrane maltotriose transport (though this can still be present) (Krogerus and Gibson, 2020). Our assessment here of the uptake of radiolabeled maltotriose demonstrates conclusively that transmembrane transport occurs across cell membranes of *S. jurei*. This was only apparent after cells had been propagated on maltotriose, indicating that the expression of the phenotype (either through gene expression or protein configuration or localization) is either limited by the presence of other sugars or requires induction by maltotriose. Known maltotriose transporters such as AGT1 and MTT1 can carry both maltose and maltotriose and do not require specific conditions for their expression (Magalhães et al., 2016). The mechanisms by which maltotriose is consumed by *S. jurei* require further investigation. Nikulin et al., in a study of brewing potential in *S. paradoxus*, noted that the efficient uptake of maltose was influenced by the growth medium, with growth on glucose leading to an extended lag phase prior to maltose use during fermentation (Nikulin et al., 2020). This lag phase was significantly reduced if yeasts were propagated on maltose. These results, and our observation that maltotriose use requires previous exposure to this sugar, suggest that brewing with *S. jurei* would benefit from carefully regulated propagation conditions.

The presence of an active trans-membrane-transport system does not exclude the possibility of *S. jurei* also hydrolyzing maltotriose extracellularly. Genes encoding both potential maltose/maltotriose transporters and extracellular α-glucosidases were found within the *S. jurei* genome. It is expected that a more thorough knowledge of the maltotriose utilization mechanisms may help to improve the potential brewing performance of *S. jurei*, both in terms of maltotriose utilization efficiency and duration of lag phase.

While brewing efficiency is a critically important trait for brewers, the contribution of yeast to beer flavor is more directly relevant for consumers. A new brewing strain should preferably offer a novel sensorial experience to beer drinkers. In this regard, the concentration of yeast-derived volatile aroma compounds is significant. The three strains included in the brewing trials had distinct flavor profiles, with each producing a high concentration of at least one important flavor-active compound. In the case of *S. jurei*, this was ethyl hexanoate, a generally desirable ethyl ester with an apple, cherry or aniseed aroma. The high concentration may have contributed to the tropical fruit notes detected by the sensory panel. In contrast to *S. jurei*, the dominant flavor volatile in the *S. pastorianus* beer was 3-methylbutylacetate. This is a highly desirable flavor compound in commercial brewing and contributes a banana or pear aroma to beer (Meilgaard, 1975). In the *S. eubayanus* beer, the typically rose-like phenylethanol was prominent. This has previously been noted for this strain (Gibson et al., 2013). Phenylethanol has previously been found to mask the perception of other flavor compounds (Bamforth, 2020) and may explain the relatively low perception of phenolic flavor notes in the *S. eubayanus* beer relative to the *S. jurei* beer.

The discovery of relatively good fermentation efficiency in *S. jurei* inspired a direct comparison with fermentation performance in other members of the strain. This comparative study included an ale *S. cerevisiae* strain, a lager *S. pastorianus* strain and six type strains of wild *Saccharomyces* species. The only species excluded was *S. uvarum*, the type strain of which was shown previously to be strongly maltose-positive, but maltotriose-negative (Nikulin et al., 2018). The species included were highly variable with respect to fermentation characteristics. Relative to the reference strains, the wild strains tended to have long lag periods following the initial fermentation of monosaccharides, and when fermentation increased, it was at a relatively low rate and often limited in extent. Given these features, it is unsurprising that wild *Saccharomyces* species might be at a competitive disadvantage compared to domesticated yeasts, and thus rarely encountered in wort fermentations. Performance varied however, not just between brewing yeasts and wild yeasts, but also amongst the wild species. As observed previously (Nikulin et al., 2018), *S. mikatae* had only limited fermentation capability, with apparently no ability to metabolize maltose. Other species appeared to be able to utilize this disaccharide, albeit at different rates and after different periods of adaptation following the initial fermentation. Of note was the high level of alcohol production by the type strain of *S. jurei*, suggesting that a superior fermentation efficiency due to maltotriose utilization is not restricted to the Bavarian strain included in the previous fermentation trials.

As well as the association between woodland habitats and yeast, there has been traditionally a strong association between wood-based materials and brewing. Wood, in addition to being the main building material for millennia, has also been used for vessels used in food production and storage. The usage of wooden materials in brewing was not only limited to vessels, it also made up tools and additives for beer production. Oak wood was used for vats, casks and barrels; chips or shavings of hornbeam (*Carpinus betulus*) and hazel (*Coryllus avellana*) have been used as clarifying aids; and spruce and birch were commonly used for making barrel bungs (Schnegg, 1921; 1922). Schnegg, in his textbook about microscopy for brewers, describes oak wood as being hard but porous with the pores being ideal for harboring microorganisms and residues of beer (Schnegg, 1921). Ash and oak trees have also been described as holy trees in Indo-Germanic culture (Dumont, 1992). Oaks have served as an inoculum for roman wine fermentations and ash trees were often tapped for their tree sap (Feier et al., 2019). Through its unique structure and capability of harboring microorganisms in nature, wood could potentially transfer yeasts from natural habitats to human made fermentations and thereby initiate the process of domestication. Due to the role of both ash and oak and their proven association with yeasts capable of fermenting cereal-based worts it is not unlikely that these trees have served as an inoculum for fermentation throughout history. While many isolates of the *Saccharomyces* species have been associated with oak trees or other members of the order of *Fagales* (containing oak trees, beeches, *Nothofagus* spp. and aforementioned *Castanopsis*), *F. excelsior* does not belong to this order. No yeasts other than *S. jurei* TUM 629 were isolated from the sampled ash tree and little is known about the specific microbiome of ash trees. Other studies have revealed evidence for the presence of *S. jurei* in different habitats (Alsammar et al., 2019). The seemingly low abundance of *S. jurei* in nature may be caused by the disadvantage of culturing methods for isolating species that are present at low numbers in the sample. However, DNA-based methods of describing complex populations (metagenomics) may also run the risk of biases in this situation (Kebschull and Zador, 2015). Recently, another strain of *S. jurei* has been isolated from the bark of an ash in the black forest, Germany (current study, data not published). More research is needed to assess the role of *S. jurei* in the ecology of yeasts, its preferred habitats and geographical distribution.

## 5 Acknowledgements

Eero Mattila and Niklas Fred are thanked for assistance in the VTT Pilot Brewery, and Aila Siltala for skilled technical assistance. Henrik Siegumfeldt (University of Copenhagen) is thanked for supervising the work of Tiina Kuusisto. Franziska Elisath (BLQ) is thanked for her work in the laboratory.

## 6 Author Contributions

TK: fermentation trials, analysis of fermentation data. FM: transmembrane transport assays. KK: whole-genome analyses. BG: first draft, conceptualization. MM: sensory and aroma analysis on the bottled beers, first draft, conceptualization. MH: first draft, isolation protocol, species identification, physiological characterization, conceptualization. OK: strain isolation, finalization of the manuscript. All authors contributed to the article and approved the submitted version.

