## Supplementary Material for "Unique brewing-relevant properties of a strain of *Saccharomyces jurei* isolated from ash (*Fraxinus excelsior*)"

### 1 Supplementary Data

*Saccharomyces jurei* TUM 629, partial ITS1-5.8S-ITS2 rDNA:

GAAGGATCATTAAAGAAATTTAATAATTTTGAAAATGGATTTTTTTGTTTTGGCAAGAGC
ATGAGCTTTTACTGGGCAAGAATACAAGAGATGGAGAGTCCAGTGGGGCCTGCGCTTAA
GTGCGCGGTCTTACTAGGCTTGTAAGTTTCTTTCTTGCTATTCCAAACAGTGAGAGATCT
TTGTGTTTTTGTTATAGGACAATTAAAACCGTTTCAATACAACACACTGTGGAGTTTTTA
TATCTTTGCAACTTTTTCTTTGGGCTTTCGAGCAATCGAGGCCCAGAGGTAACAAACACA
AACAATTTTATTTATTCATTAAATTTTTGTCAAAAACAAGAATTTTCGTAACTGGAAATT
TTAAAAATATTAAAAACTTTCAACAACGGATCTCTTGGTTCTCGCATCGATGAAGAACG
CAGCGAAATGCGATACGTAATGTGAATTGCAGAATTCCGTGAATCATCGAATCTTTGAA
CGCACATTGCGCCCCTTGGTATTCCAGGGGGCATGCCTGTTTGAGCGTCATTTCCTTCTC
AAACATTCTGTTTGGTAGTGAGTGATACTCTTTGGAGTTAACTTGAAATTGCTGGCCTTT
TCATTGGATGTTTTTTTCCAAAGAGAGGTTTCTCTGCGTGCTTGAGGTATAATGCAAGTA
CGGTCGTTTTAGGTTTTACCAACTGCGGCTAATCTTTTTTGTACTGAGCGTATTGGAACG
TTATCGATAAGAAGAGAGCGTCTAGGCGAACAATGTTCTTAAAGTTGACCTCAA

*Saccharomyces jurei* TUM 629, partial D1/D2 26S rDNA:

TAAGCAAGCATCCTTGACTTACGTCGCAGTCCTCAGTCCCAGCTGGCAGTATTCCCGCAG
GCTATAATACTTCCCGAAGCAAGTTACATTCCTACGGATTTATCCTGCCACCAAAACTGA
TGCTGGCCCAGTGAAATGCGAGATTCCCCTACCCACAAGGAGCAGAGGGCACAAAACA
CCATGTCTGATCAAATGCCCTTCCCTTTCAACAATTTCACGTACTTTTTCACTCTCTTTTC
AAAGTTCTTTTCATCTTTCCATCACTGTACTTGTTCGCTATCGGTCTCTCGCCAATATTTA
GCTTTAGATGGAATTTACCACCCACTTAGAGCTGCATTCCCAAACAACTCGACTCTTCGA
AGGCACTTTACATAGAACCGCACTCCTCGCCACACGGGATTCTCACCCTCTATGACGTCC
TGTTCCAAGGAACATAGACAAGGAACGGCCCCAAAGTTGCCCTCTCCAAATTACAACTC
GGGCACCAAAGGTACCAGATTTCAAATTTGAGCTTTTGCCGCTTCACTCGCCGTTACTAA
GGCAATCCCGGTTGGTTTCTTTTCCTCCGCTTATTG

### 2 Supplementary Tables

**Supplementary Table S1**: Genes in S. jurei NCYC 3947$^{T}$ (accession number GCA_900290405) encoding potential maltotriose-transporting permeases

| Gene | Chromosome | Start | End | Identity to S. cerevisiae S288C (%) |
|---|---|---|---|---|
| *MAL31*-like | LT986468.1 | 16347 | 18189 | 84,1 % |
| *MAL31*-like | LT986469.1 | 706843 | 704987 | 82,2 % |
| *MAL31*-like | LT986469.1 | 719396 | 717547 | 84,4 % |
| *MAL11*-like | LT986469.1 | 695682 | 693841 | 82,6 % |
| *IMA5*-like | LT986469.1 | 32730 | 31005 | 78,9 % |
| *IMA5*-like | LT986472.1 | 16533 | 14788 | 84,0 % |
| *IMA5*-like | LT986472.1 | 724412 | 726157 | 84,0 % |

### 3 Supplementary Figures

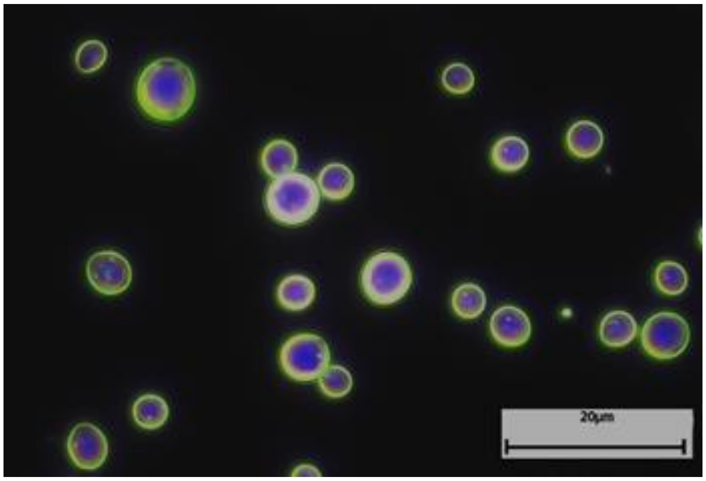


**Supplementary Figure 1.** Cell morphology of *S. jurei* TUM 629 (48 h liquid culture in 12.4 °P Brewer's wort from pale barley malt); dark field microscopy microscope (Axilob 5, Zeiss GmbH, Germany).

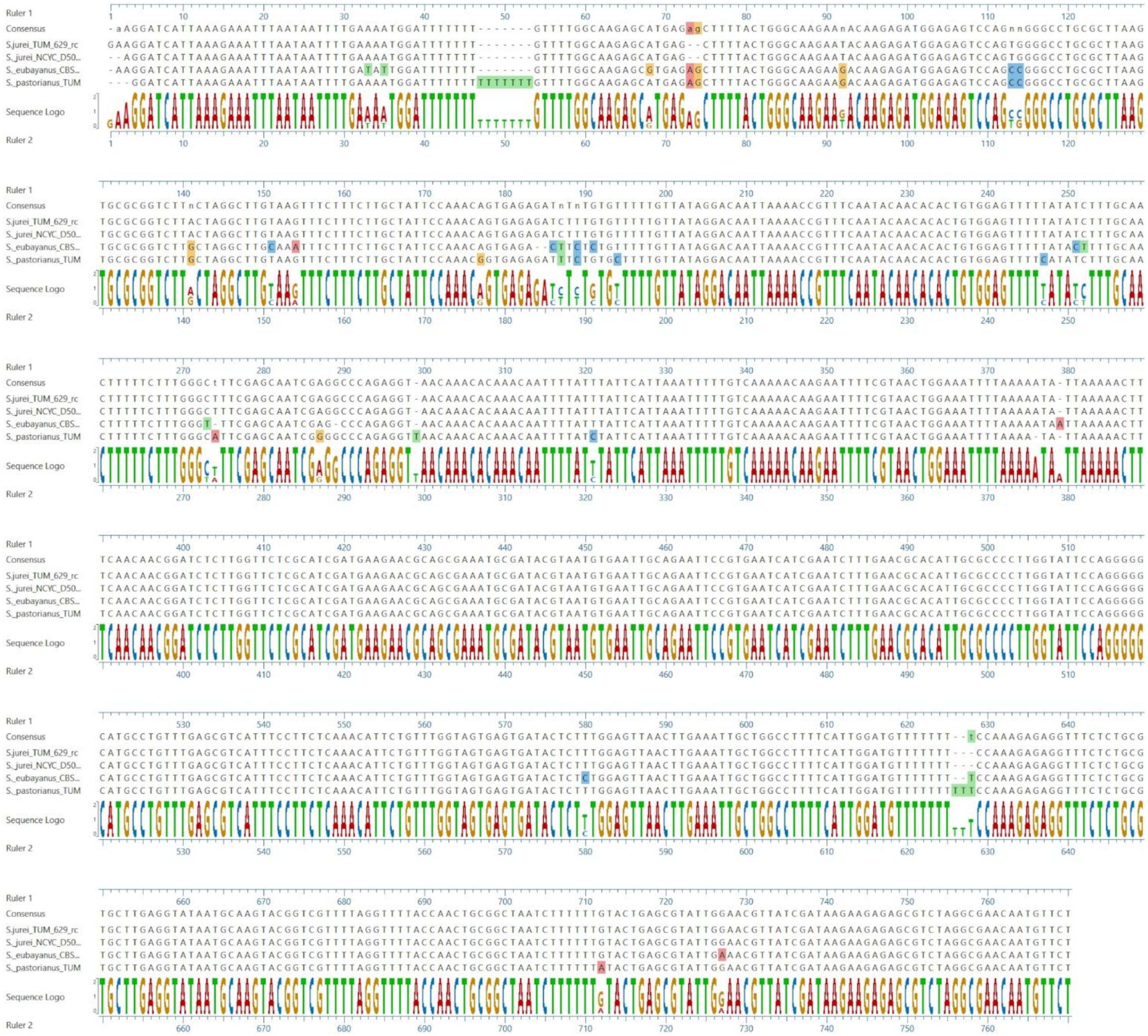


**Supplementary Figure 2.** Clustal W Alignment (multi sequence alignment) of *S. jurei* type strain NCYC 3947$^{T}$, TUM 629, *S. eubayanus* C902 (CBS 12357$^{T}$) and *S. pastorianus* TUM 34/70.

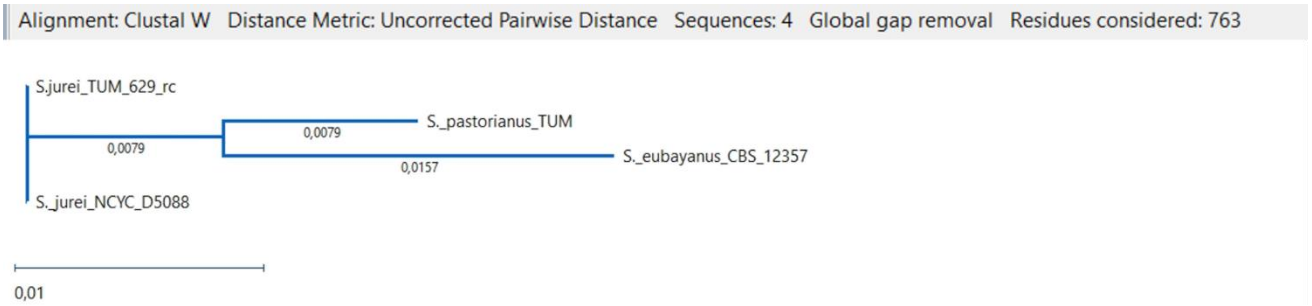


**Supplementary Figure 3.** Phylogenetic tree resulting from the Clustal W Alignment of *S. jurei* type strain NCYC 3947$^{T}$, TUM 629, *S. eubayanus* C902 (CBS 12357$^{T}$) and *S. pastorianus* TUM 34/70 showing the species identity of the newly isolated *S. jurei* strain.

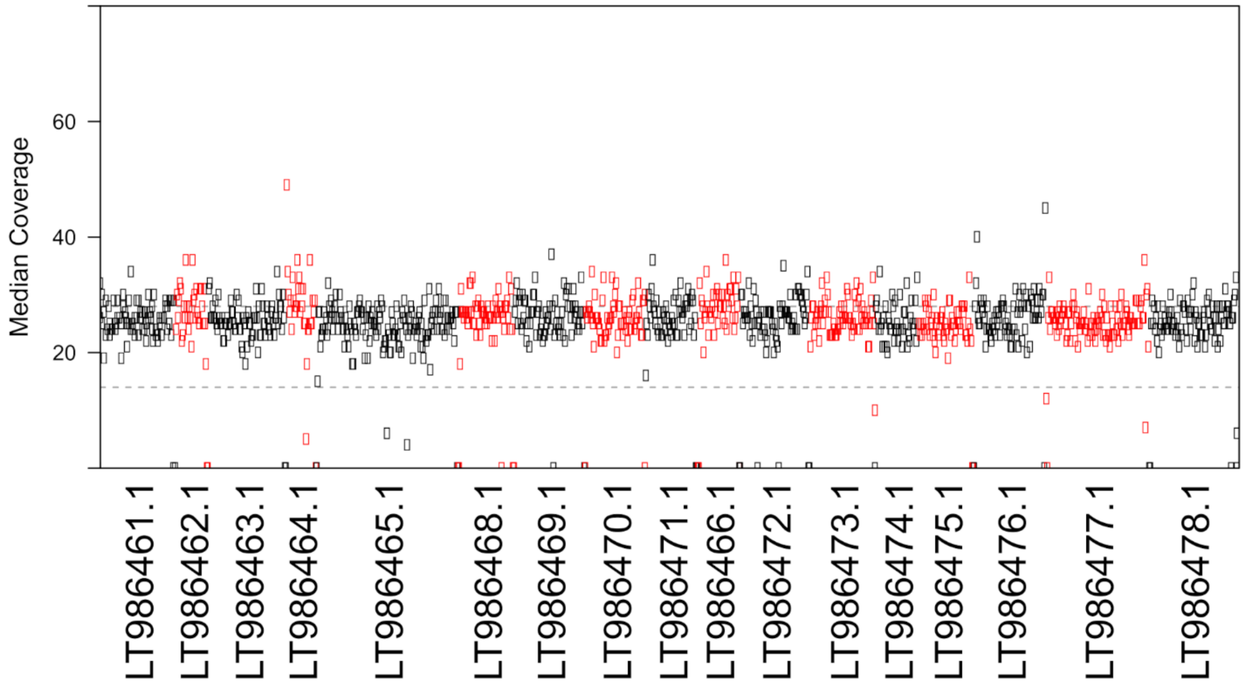


**Supplementary Figure 4.** The sequencing coverage (median coverage in 10 kbp windows) of reads from *S. jurei* TUM 629 aligned to a *S. jurei* NCYC 3947$^{T}$ (NCBI accession number GCA_900290405; (Naseeb et al., 2018)) reference genome.

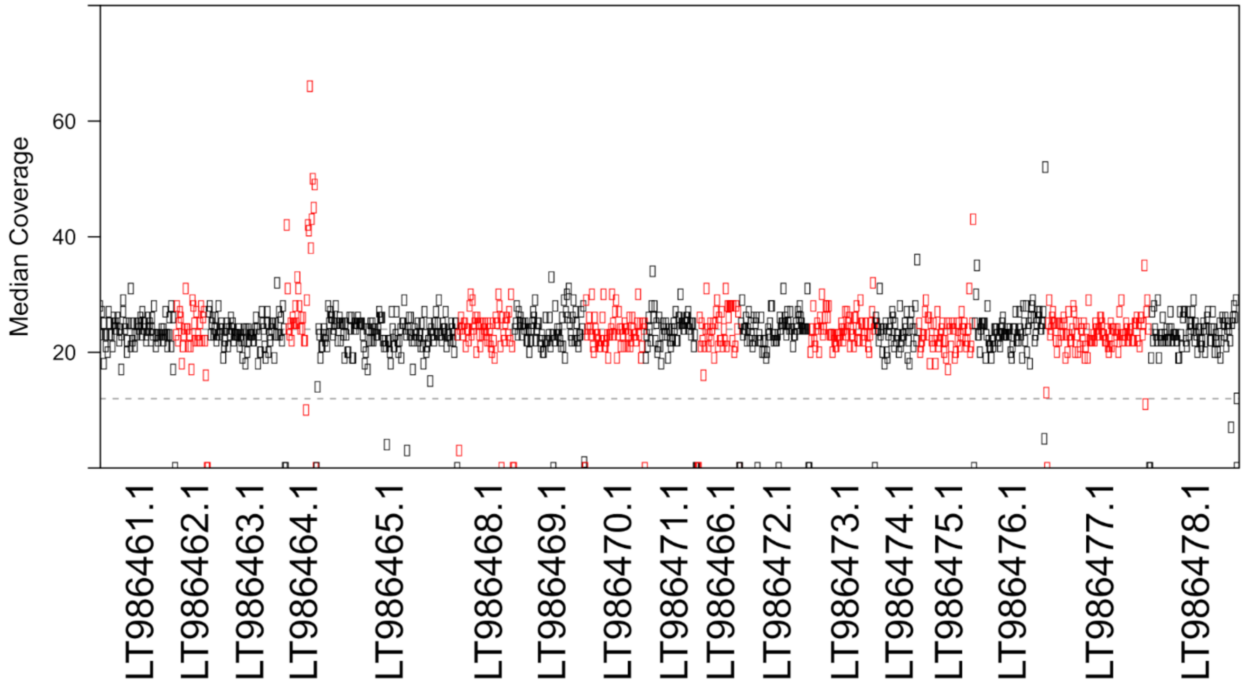

**Supplementary Figure 5.** The sequencing coverage (median coverage in 10 kbp windows) of reads from *S. jurei* VTT C-171003$^T$ aligned to a *S. jurei* NCYC 3947$^T$ (NCBI accession number GCA_900290405; (Naseeb et al., 2018)) reference genome.
